# DNA barcoding and species delimitation based on four Chloroplast loci to identify species within the Elaeocarpaceae family

**DOI:** 10.1101/2024.03.14.585135

**Authors:** Jyotsana Khushwaha, Alpana Joshi, Subrata K. Das

## Abstract

DNA barcoding is an indispensable taxonomic tool for plant species identification. The present research successfully amplified the four coding regions of plastid namely; *matK*, *rpoB*, *ndhJ*, and *accD*, sequenced, and submitted it to NCBI Genbank after verification. This study evaluated the significance of single (*matK*, *rpoB*, *ndhJ*, and *accD*) and two-locus plastid loci (*rpoB+matK*, *ndhJ*+*matK*, *accD*+*matK*) and examined their abilities for species identification and phylogenetic construction of Elaeocarpaceae plants. The average aligned length for *matK*, *rpoB*, *ndhJ*, *accD*, *ndhJ*+*matK*, *rpoB*+*matK*, and *accD*+*matK* was 395 bp, 538 bp, 477 bp, 254 bp, 872 bp, 937 bp, and 649 bp, respectively. The highest substitution rate was recorded in the *rpoB* (18.47%), followed by *ndhJ* (17.38%), *accD* (14.00%), and *matK* (11.23%). The number of variable sites, segregating sites (S), nucleotide diversity (Pi), and mutations (Eta) were observed to be highest in the *matK* region. Evolutionary relationship based on the Neighbor-joining method demonstrated that candidate barcode sequences were competent for identifying *Crinodendron*, *Elaeocarpus*, *Sloanea*, and *Vallea* species. A significant barcoding gap was identified in two barcoding regions (*matK* and *ndhJ*+*matK*). The species-specific barcodes for the genus *Elaeocarpus* were generated based on Single Nucleotide Polymorphism (SNP) analysis. This study offers novel strategies for effectively identifying species within Elaeocarpaceae and lays the foundation for germplasm conservation and utilization.

## Introduction

Elaeocarpaceae is a huge family comprising more than 550 known species distributed in 12 genera and represented mainly in Australia, South America, and Southeast Asia [1,2]. Genus *Elaeocarpus* comprises 350-400 species dispersed in lowland to montane areas of Asia, Australia, Madagascar, and the Pacific islands [3,4,5]. The *Crinodendron*, *Sloanea,* and *Vallea* are endemic genera in South America, whereas nine other genera are distributed in Australia [1,2].

Phylogenetic studies on Elaeocarpaceae have been mainly conducted based on morphological features [3,6,7,8]. However, a few studies were performed based on fruit mesocarp morphology because fruit mesocarp displays significant variation compared to other morphological characteristics [9,10]. DNA barcoding is an integrated taxonomic tool to identify and distinguish species using short-standardized sequences. Recently, DNA barcoding has emerged as a vital taxonomic tool due to its simplicity, accuracy, repeatability, and efficiency [11]. Mitochondrial *CO1* (cytochrome c oxidase subunits I) was proposed as a universal barcode in plants and successfully applied to species identification of animals, birds, fish, insects, and nematodes [12,13,14]. However, due to its slow evolution, *COI* is not considered a suitable barcode for the plant kingdom. Among the several barcode regions tested by various researchers, the Consortium for the Barcode of Life (COBL) recommended *rbcL* and *matK* loci as preferred barcode loci for identifying plant species [15]. Finding an appropriate DNA barcode for plant species identification is challenging because the ideal barcodes should have high inter- and low intra-specific divergence. High taxonomic coverage and better resolution capacity are critical features of an ideal DNA barcode.

Plastid DNA barcode (*atpF-atpH* (ATP synthase subunit CFO I-ATP synthase subunit CFO III), *accD* (Acetyl-CoA Carboxylase), *matK* (Maturase K), *ndhJ* (NADH Dehydrogenase J Subunit), *psbK-psbI* (photosystem II protein K-photosystem II protein I), *rpoB* (RNA Polymerase subunits B), *trnH-psbA* (transfer RNA Histidine-photosystem II protein D1), and *ycf1b* (TIC complex) and nuclear DNA barcode regions (Internal Transcribed Spacer (*ITS*) are commonly applied in DNA barcoding of higher plants [16,17,18,19]. Several studies have investigated the Phylogenetic relationship within Elaeocarpaceae based on nuclear and plastid DNA sequences [1,20,21]. These studies focused on determining evolutionary relationships using nuclear *ITS* and plastid *trnL-trnF* sequences within Elaeocarpaceae. Another study used *ITS* and *trnL-trnF* sequence data of 13 *Elaeocarpus* species to establish intergeneric relationships within Elaeocarpaceae (mainly from Australia) [1]. The evolutionary studies using *Elaeocarpus* taxa were performed based on *ITS* sequence data [20]. Similarly, phylogenetic relationships of 59 Elaeocarpus taxa were established using plastid (*trnL-trnF*, *trnV-ndhC* intergenic spacer) and nuclear (*ITS*) sequences [7,8]. Collectively, these molecular analyses had satisfactorily resolved the evolutionary relationships between the majority of Elaeocarpaceae genera. The evolutionary relationships among most of the *Elaeocarpus* species are not yet established because of low sequence divergence, although some clades were determined in several phylogenetic studies [1,7,8,19,20,21]. However, various research groups have suggested different combinations of these loci as potential DNA barcodes, but until now, no barcode can discriminate plants at the species level. Single Nucleotide Polymorphisms are among the best measures to distinguish closely related species. Specific barcodes based on Single Nucleotide Polymorphism have been generated to identify particular plant species [18,22].

In the present study, we used four chloroplast-based sequences (*matK*, *rpoB, ndhJ*, *accD*) and three combined sequences, including *ndhJ*+*matK*, *rpoB*+*matK*, *accD*+*matK* to develop species-specific barcode of Elaeocarpaceae plants based on Neighbor-joining (NJ) method and Single Nucleotide Polymorphism (SNP) analysis. This study provides an innovative method of developing species-specific DNA barcodes of Elaeocarpaceae plants and establishes the foundation for preserving, assessing, and utilizing Elaeocarpaceae germplasms.

## Materials and Methods

### DNA Amplification and Sequencing

High-quality genomic DNA was extracted from the young leaf samples of *E. ganitrus* using DNeasy Plant Mini Kit (Qiagen, Germany) per the manufacturer’s instructions [19]. PCR-based amplification of four barcoding regions was performed using *matK* (*matK*_F: 5’-CGATCTATTCATTCAATATTTC 3’ and *matK*_R: 5’-TCTAGCACACGAAAGTCGAAGT-3’), *ndhJ* (*ndhJ*_F: 5’-TTGGGCTTCGATTACCAAGG-3’ and *ndhJ*_R: 5’-TCAATGAGCATCTTGTATTTC-3’), *rpoB* (*rpoB*_F: 5’-ATGCAACGTCAAGCAGTTCC-3’ and *rpoB*_F: 5’-GATCCCAGCATCACAATTCC-3’), and *accD* (*accD*_F: 5’-AGTATGGGATCCGTAGTAGG -3’ and *accD*_F: 5’-TCTTTTACCCGCAAATGCAAT-3’). The PCR thermal profile included initial denaturation of 94 °C for 4 min, followed by 35 cycles of denaturation at 94 °C for 30 s, primer annealing at 55 °C for 40 s, and extension at 72 °C for 1 min, with a final extension of 10 min at 72 °C. The PCR-amplified products were examined via electrophoresis in a 1.2% agarose gel. The purified product was used for sequencing reaction per the kit’s protocol (BigDye Terminator v3.1, Applied Biosystems).

### Nucleotide Sequences

In the present study, 99 *matK*, 36 *rpoB*, 34 *ndhJ*, and 34 *accD* sequences of the Elaeocarpaceae family were retrieved from the NCBI database (https://www.ncbi.nlm.nih.gov/) for further analyses. All retrieved sequences were subjected to a critical evaluation, and low-quality and short sequences were removed, like sequences containing <3% ambiguous base ’N’ [23]. The accession numbers included in this study are provided in Table 1. The combined sequence, including *rpoB*[+[*matK*, *ndhJ*+*matK*, and *accD*+*matK*, was obtained by the supermat function in the R Phylotools package [18].

### Sequence Alignment and Data Analysis

The sequence alignment for each locus (*accD*, *matK*, *ndhJ*, *rpoB*, *ndhJ*+*matK*, *rpoB*+*matK*, *accD*+*matK*) was performed using the MUSCLE algorithm of MEGA v11 [24]. The number of INDELs, polymorphic site, and neutrality tests (Fu & Li’s D, Fu & Li’s F, and Tajima’s D) were performed using the DnaSP v6.12 (URL: http://www.ub.edu/dnasp/index_v5.html) [18,19,25]. Phylogenetic trees were constructed with the Neighbor-joining (NJ) method with 10,000 bootstraps using the Kimura 2-Parameter (K2P) model in MEGA v11 [26].

### Analysis of Barcoding gap

Barcoding gap analysis of each locus (*accD*, *matK*, *ndhJ*, *rpoB*, *ndhJ*+*matK*, *rpoB*+*matK*, *accD*+*matK*) was conducted using Automatic Barcode Gap Discovery (ABGD) (https://bioinfo.mnhn.fr/abi/public/abgd/abgdweb.html) and Assemble Species by Automatic Partitioning (ASAP) (https://bioinfo.mnhn.fr/abi/public/asap/) web servers [26,27]. A two-dimensional QR (Quick Response) code was generated for a specific candidate barcode marker(s) using the QR Code Web Server (https://products.aspose.app/barcode/generate).

## Results and Discussion

### Amplification and Sequencing of Chloroplast Sequences

The chloroplast sequences of four different coding regions such as *matK* (Maturase K), *ndhJ* (NADH Dehydrogenase J Subunit), *rpoB* (RNA Polymerase subunits B), and *accD* (Acetyl-CoA Carboxylase) were amplified using gene-specific primers for each barcode. The amplification of the *matK* gene using the *matK*_F and *matK*_R primer pair produced the desired 460 bp fragment. The *ndhJ* gene has been amplified using the *ndhJ*_F and *ndhJ*_R primer pair, and an amplicon of the desired length (422 bp) was recorded. The *accD* gene has been amplified using the *accD*_F and *accD*_R primer pair, and an amplicon of 299 bp was obtained. The *rpoB* gene has been amplified using the *rpoB*_F and *rpoB*_R primer pair and produced the desired fragment of 542 bp in size (Fig. 1). The sequence of each barcode was compared using the NCBI blast tool for similarity search. The top ten BLASTn scores for each barcode are presented in Table 1. Sequences were aligned using Bioedit software, and the final contigs of each sequence were deposited in the National Center for Biotechnology (NCBI) GenBank to obtain accession numbers: OR073889 (*rpoB*), OR073890 (*accD*), and OR073891 (*ndhJ*).

**Fig. 1.**
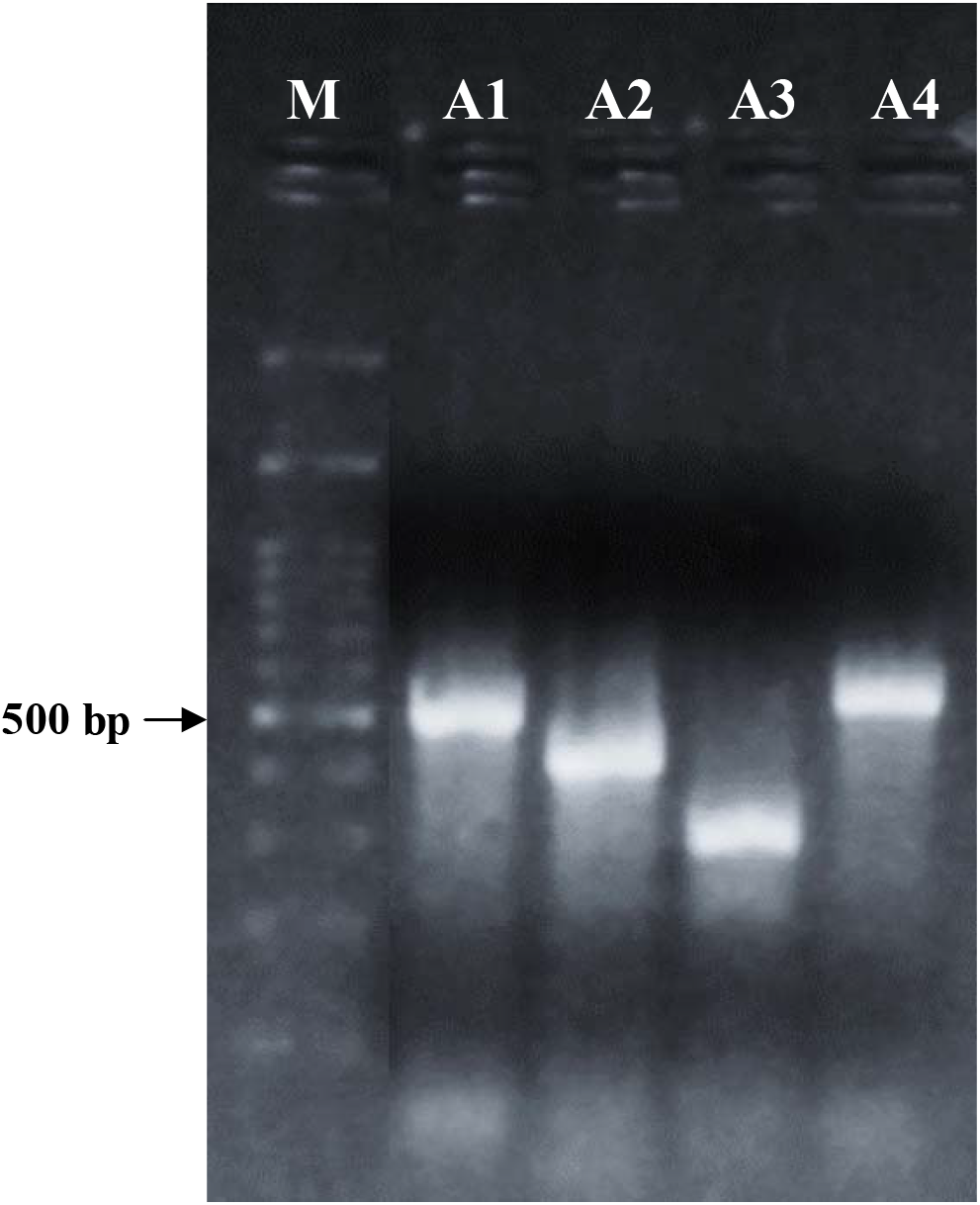
Agarose gel electrophoresis (1.2%) showing PCR-amplified band of four different barcoding regions. **M:** 1 kb plus DNA ladder, **A1:** *matK* (460bp), **A2:** *ndhJ* (422bp), **A3:** *accD* (299bp), **A4:** *rpoB* (542bp).

### Nucleotide Sequence analysis

In this study, 99 *matK*, 36 *rpoB*, 34 *ndhJ*, and 34 *accD* of the Elaeocarpaceae family were collected from the NCBI database (https://www.ncbi.nlm.nih.gov/) for further analyses. After blasting and editing, the aligned length of *matK*, *rpoB*, *ndhJ*, and *accD* was 395 bp, 538 bp, 477 bp, and 254 bp, respectively, and that of combined sequence including *ndhJ*+*matK*, *rpoB*+*matK*, *accD*+*matK* were 872 bp, 937 bp, 649 bp, respectively.

The nucleotide frequencies of *matK* for T, C, A, and G were 33.58 %, 15.98%, 34.12 %, and 16.29 %, respectively, and the average GC contents ranged from 28.86 % to 34.14 %. The nucleotide frequencies of *rpoB* for T, C, A, and G were 30.32 %, 23.56 %, 26.19 %, and 19.91 %, respectively, and the average GC contents varied from 35.34 % to 51.02 %. The nucleotide frequencies of *ndhJ* for T, C, A, and G were 28.00 %, 23.76 %, 31.36 %, and 16.94 %, respectively, and the average GC contents ranged from 33.02 % to 50.90 %. The nucleotide frequencies of *accD* for T, C, A, and G were 33.15 %, 17.32 %, 28.28 %, and 21.23 %, respectively, and the average GC contents varied from 29.37 % to 44.70 %. The highest GC content was recorded in *rpoB*, followed by *ndhJ*, *accD*, and *matK* (Table 2).

This study assesses the substitution of the different bases in the studied barcoding regions on entire codon positions under the Tamura (1992) model [28]. The average AT and GC content of chloroplast barcodes at different coding positions of codons are displayed in Fig. 2. Two chloroplasts loci, namely *rpoB,* and *ndhJ* showed a higher transitional substitution rate than *matK* and *accD* (Table 3). The highest substitution rate from T to C and A to G was noted in the *rpoB* region (18.47%) followed by *ndhJ* (17.38%), *accD* (14.00%) *matK* (11.23%) regions. In contrast, the changing frequency from G to A and C to T of *ndhJ* (25.32%) was the highest, followed by *rpoB* (24.01%), *matK* (23.55%), and *accD* (22.30%).

**Fig. 2.**
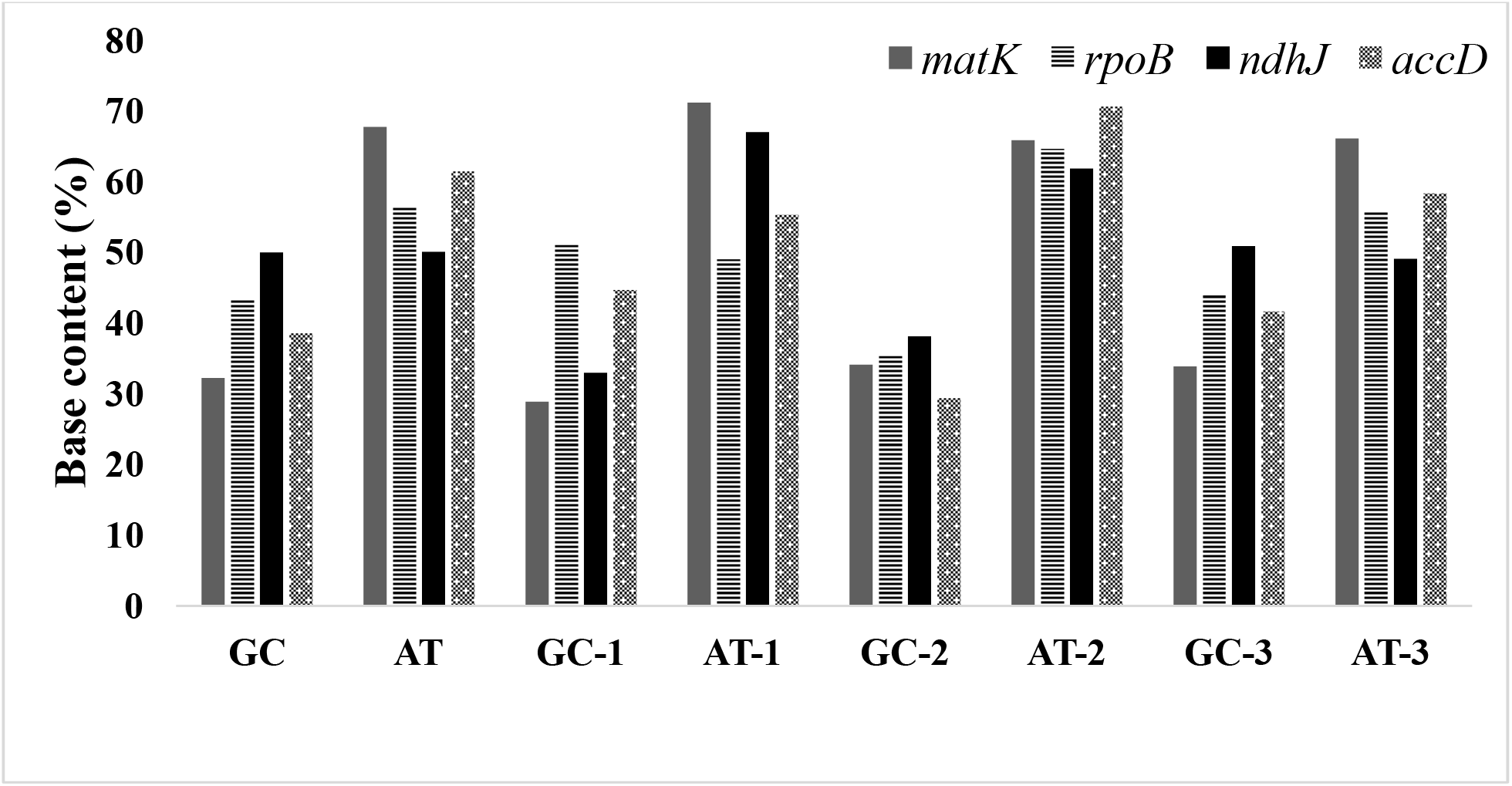
Average AT and GC content of candidate barcode sequences in Elaeocarpaceae plants.

Polymorphism site analysis of the four candidate barcodes (*matK*, *rpoB*, *ndhJ*, *accD*) and three combined barcode sequences (*ndhJ*+*matK*, *rpoB*+*matK*, *accD*+*matK*) was conducted using DnaSP v6.12 [19, 29]. Among the single loci, the *matK* sequence had the highest proportion of variable sites and parsimony informative sites, followed by *rpoB*, *ndhJ,* and *accD*. In combined sequences, *rpoB+matK* had the highest proportion of variable sites and parsimony informative sites, followed by *ndhJ*+*matK* and *accD*+*matK* (Table 4).

### Genetic diversity

The highest level of nucleotide diversity (0.01846) was recorded in *matK* sequence with the maximum θ (0.0231), S (39), k (6.48), and Eta (42) values among single gene sequences. The *rpoB*+*matK* sequences had the maximum level of nucleotide diversity (0.01833) was recorded in *matK* sequence with the maximum θ (0.0202), S (68), k (16.44), Eta (70), and Hd (0.946) values among combination gene sequences (Table 5). Moreover, the test statistics of neutrality Tajima’s D, Fu & Li’s D, and Fu & Li’s F were negative and statistically insignificant (P[>[0.02). The findings indicated that these populations were constant and less vulnerable to environmental or other external factors [30].

Population size changes were studied to determine nucleotide mismatch distribution across the Elaeocarpaceae family (Fig. 3). The maximum coefficient of variation (1.0461) was recorded in *matK* followed by *ndhJ* (0.8893), *accD* (0.800), *rpoB* (0.7936), *ndhJ+matK* (0.7864), *rpoB+matK* (0.7833), and *accD+matK* (0.7739) sequences. The mismatch distribution analysis revealed a low level of genetic diversity in Elaeocarpaceae plants. In addition, the distribution of pairwise differences was also calculated using Ramos-Onsins and Rozas statistics (R2). Among single and combined sequences, the smallest positive value of R2 statistics was recorded in *matK* (0.0810) followed by *ndhJ* (0.0830), *accD* (0.0909), *rpoB* (0.1066), *ndhJ*+*matK* (0.1073), *rpoB*+*matK* (0.1087), and *accD*+*matK* (0.1163) sequences. The result indicated that *matK* gene sequences could be suitable for identifying Elaeocarpaceae plants.

**Fig. 3.**
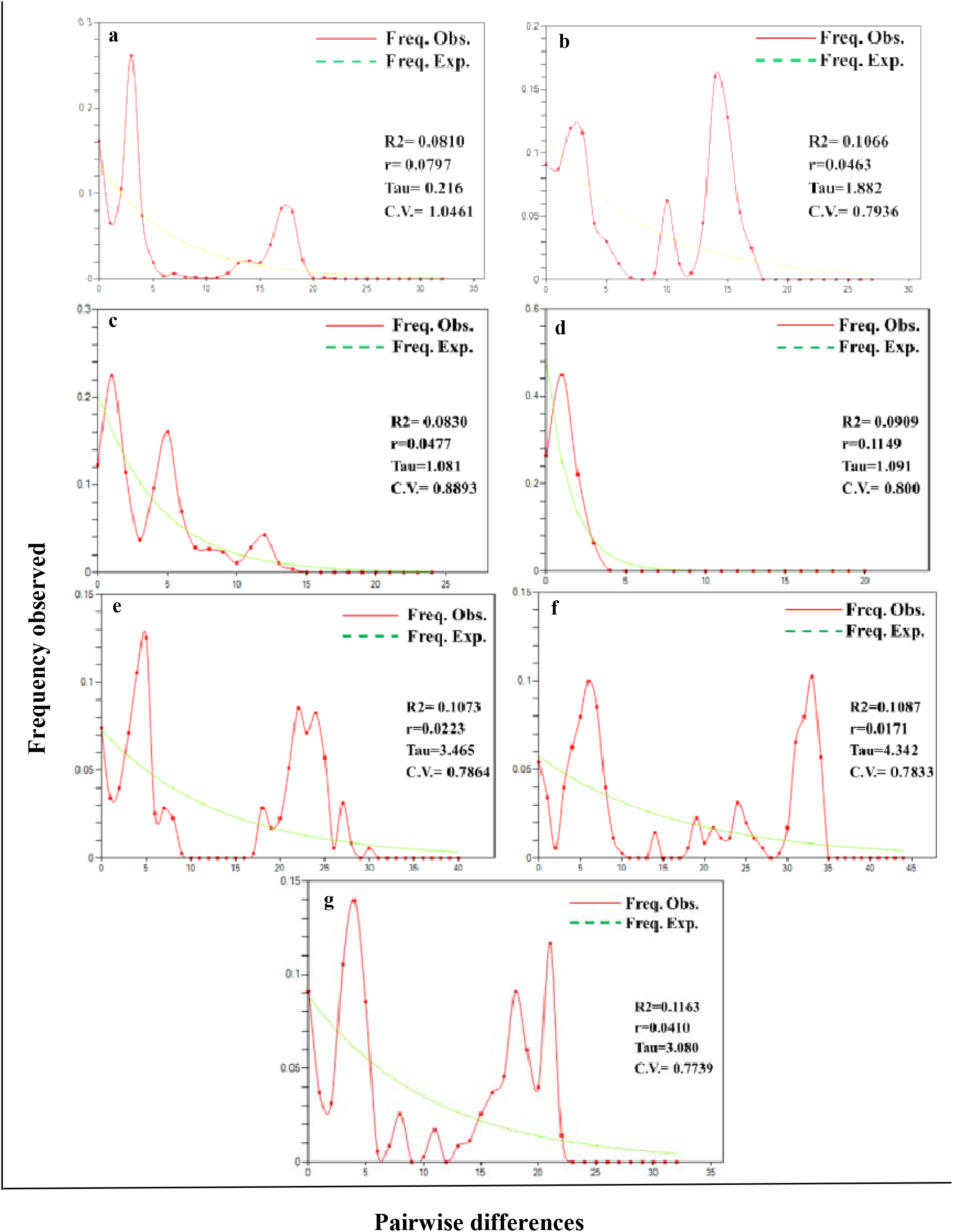
Distribution of pairwise genetic variation based on four candidate barcodes: (a) *matK*, (b) rboB, (c) *ndhJ*, (d) *accD* and three combined sequences; (e) *ndhJ*+*matK*, (f) *rpoB*+*matK*, (g) *accD*+*matK* sequences. CV: Coefficient of variation, R2: Ramos-Onsins and Rozas statistics, r: Raggedness statistic, Tau: Date of the Growth or Decline measured of mutational time.

### Phylogenetic analysis based on Single locus barcode

The final aligned data matrix of the *matK* gene contained 99 sequences and 395 nucleotides, of which 312 characters were constant, 39 characters were variable, 29 nucleotide positions were parsimony informative, and 10 were singleton. In the constructed phylogenetic tree, all the species of the Elaeocarpus were grouped with 97% bootstrap support. Two accessions of the genus *Crinodendron* (*C. patagua*; AY935929 and NC063646) clustered together with 99% bootstrap support. The genus *Sloanea* grouped with related species in a single clade with 94% bootstrap support except for one accession (*S. sinensis*, MT982367) grouped with genus *Elaeocarpus* (Fig. 4). Phylogenetic analysis based on 36 sequences (538 nucleotides) of *rpoB* gene sequence showed 326 conserved, 25 variable, 17 parsimony-informative, and eight singleton sites. An *Elaeocarpus* clade (92% bootstrap support) consisted of most taxa from the genus *Elaeocarpus* and one from the *Sloanea* (*S. sinensis*, MT982367). A similar clustering pattern was also recorded in *matK* phylogeny. Two species of the genus *Vallea* (*V. stipularis*; MW218468 and NC063647) were grouped with 99% bootstrap support. Out of ten, nine accessions of the genus *Sloanea* clustered together at the same clade with 93% bootstrap support (Fig. 5). The final aligned length of the *ndhJ* gene contained 34 sequences and 477 nucleotides, of which 452 characters were constant, 23 characters were variable, 14 nucleotide positions were parsimony informative, and 9 were singleton: most *Elaeocarpus* and one *Sloanea* species (*S. sinensis*, MT982367) clustered together at the same clade with 78% bootstrap support. The rest of the taxa of the genus Sloanea were distributed into two groups with strong bootstrap support: *S. dasycarpa*, *S. cordifolia*, and *S. sinensis,* in one group with 84% bootstrap support. Another group contains taxa *S. longiaculeata*, *S. hemsleyana*, *S. leptocarpa,* and *S. sinensis* clustered together with 60% bootstrap support. Both species of genus *Vallea* (*V. stipularis*; MW218468 and NC063647) grouped with 99% bootstrap support (Fig. 6). The phylogenetic analysis based on 34 sequences (254 nucleotides) of the *accD* gene sequence showed 247 conserved sites, seven variable sites, three parsimony-informative sites, and four singletons. The result indicated that eight taxa of the genus *Elaeocarpus* (*E. glabripetalus*, *E. braceanus*, *E. decipiens, E. sylvestris*, *E. duclouxii*, *E. serratus*, and *E. ganitrus*) resolved with 64% bootstrap, in contrast other *Elaeocarpus* taxa did not resolve in the *accD* phylogeny. Most of the species of the genus *Sloanea* clustered together in the same group with 63% bootstrap support except *S. sinensis* (MT982367), whereas both species of genus *Vallea* (*V. stipularis*; MW218468 and NC063647) grouped with 63% bootstrap support (Fig. 7). The notable observation in the phylogenetic clusters of the *accD* sequence was that the *E. arnhemicus* (MW218461) was distantly away from all closely related species.

**Fig. 4.**
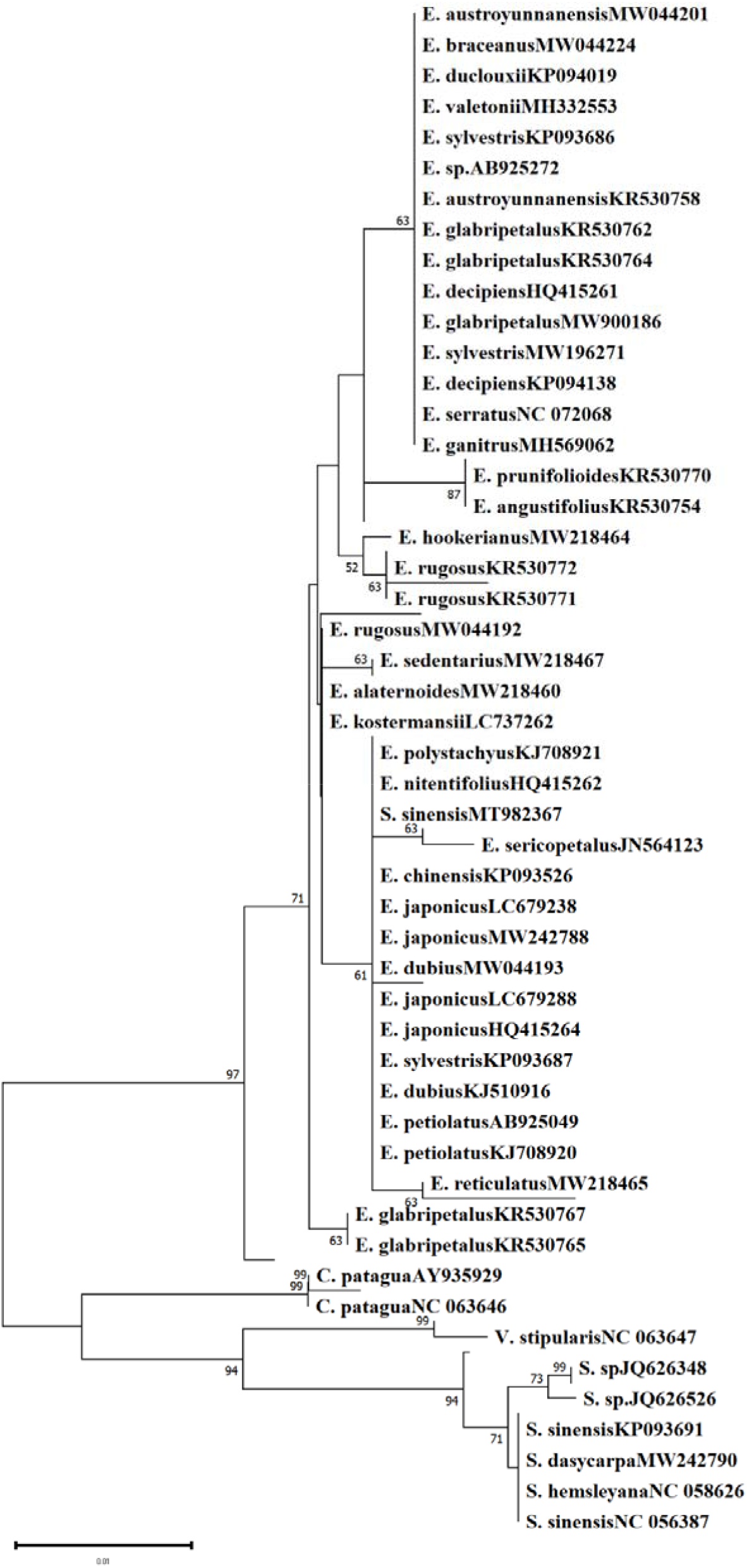
Neighbor-joining tree of Elaeocarpaceae based on *matK* sequences using Tamura 3-parameter method. This analysis involved 99 nucleotide sequences and 395 sites. The Number on the branches represent more than or equal to 50 percent support after the 10,000 bootstrap replications test.

**Fig. 5.**
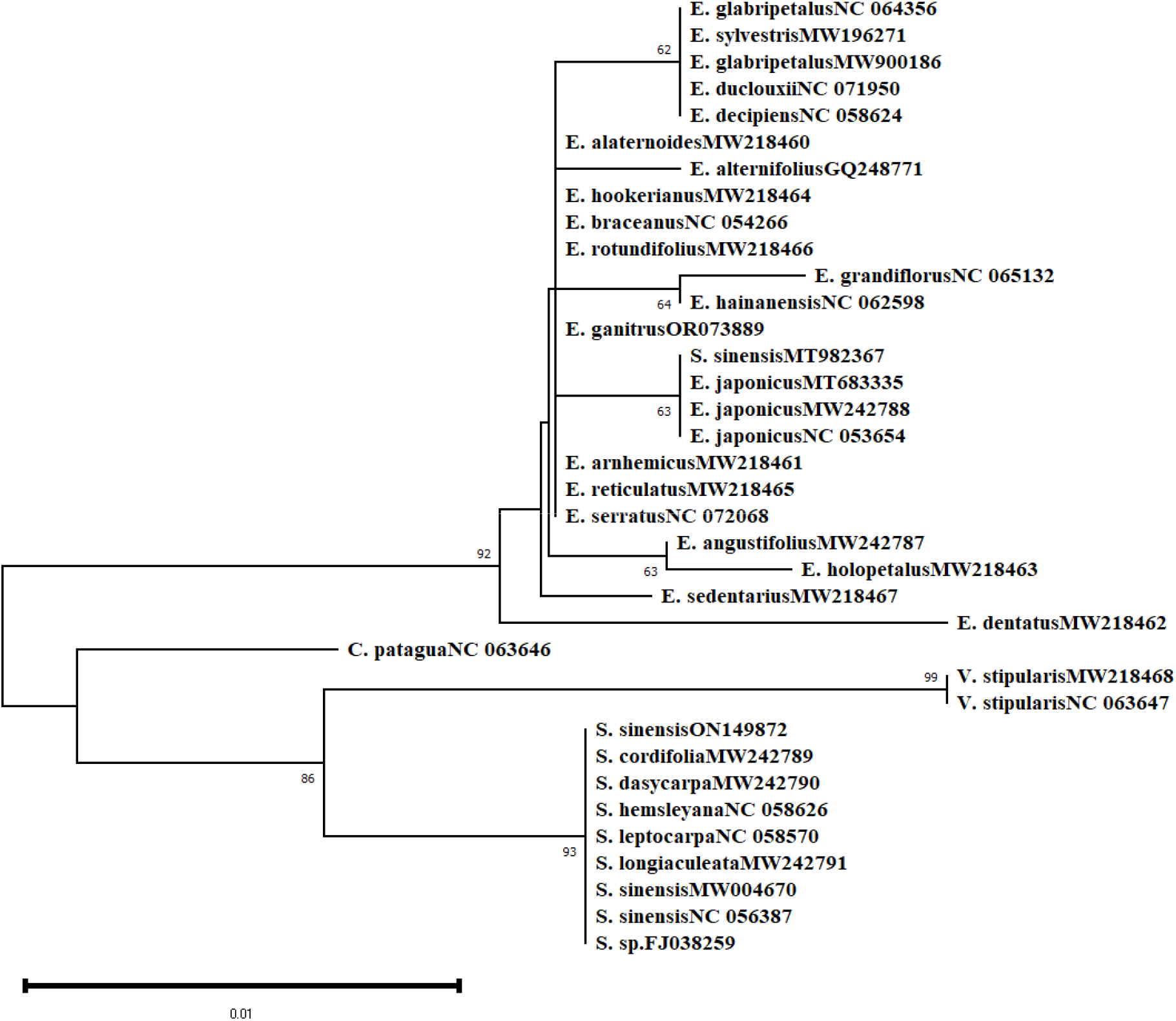
Neighbor-joining tree of Elaeocarpaceae based on *rpoB* sequences using Tamura 3-parameter method. This analysis involved 36 nucleotide sequences and 538 sites. The Number on the branches represent more than or equal to 50 percent support after the 10,000 bootstrap replications test.

**Fig. 6.**
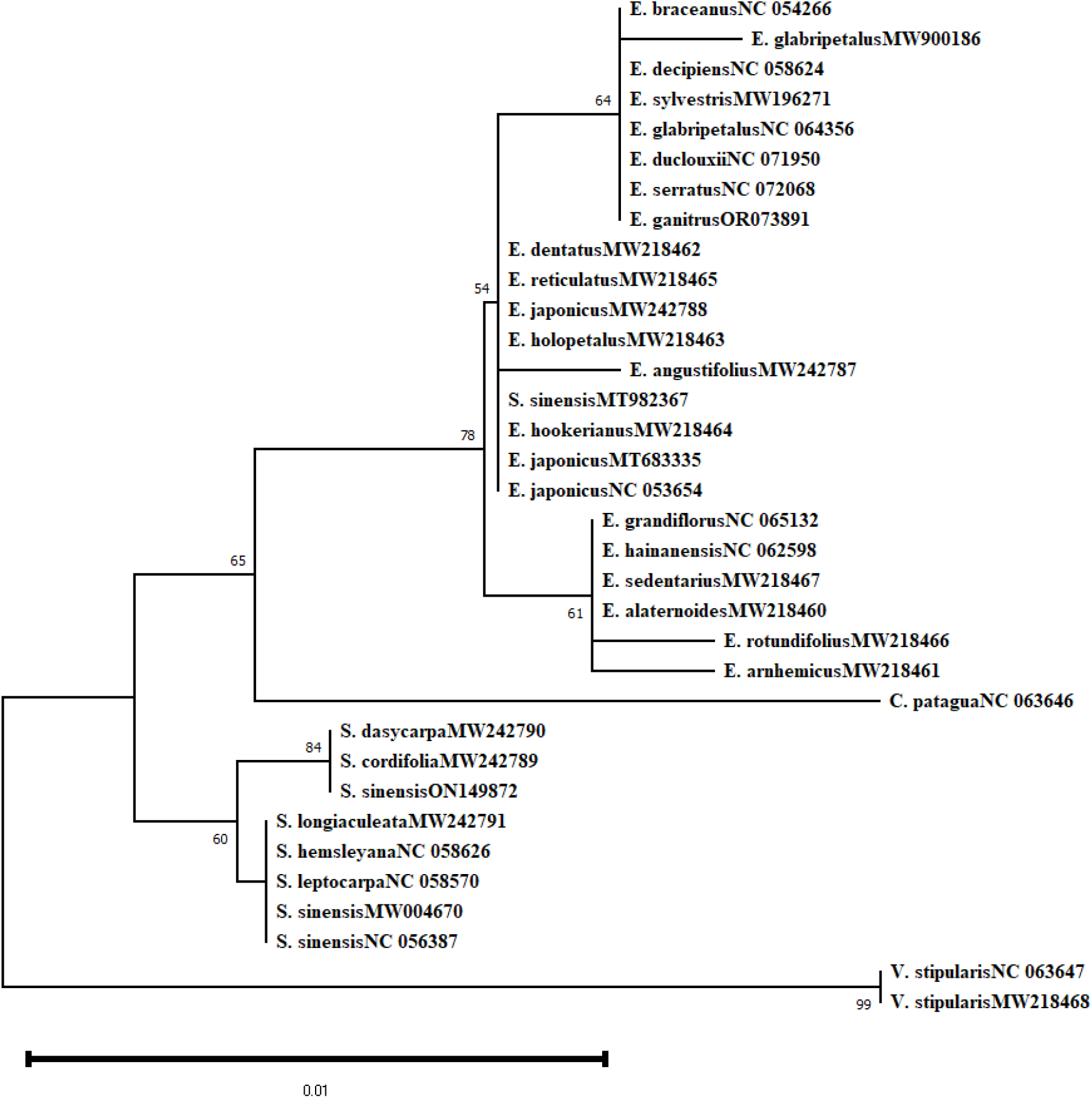
Neighbor-joining tree of Elaeocarpaceae based on *ndhJ* sequences using Tamura 3-parameter method. This analysis involved 34 nucleotide sequences and 477 sites. The Number on the branches represent more than or equal to 50 percent support after the 10,000 bootstrap replications test..

**Fig. 7.**
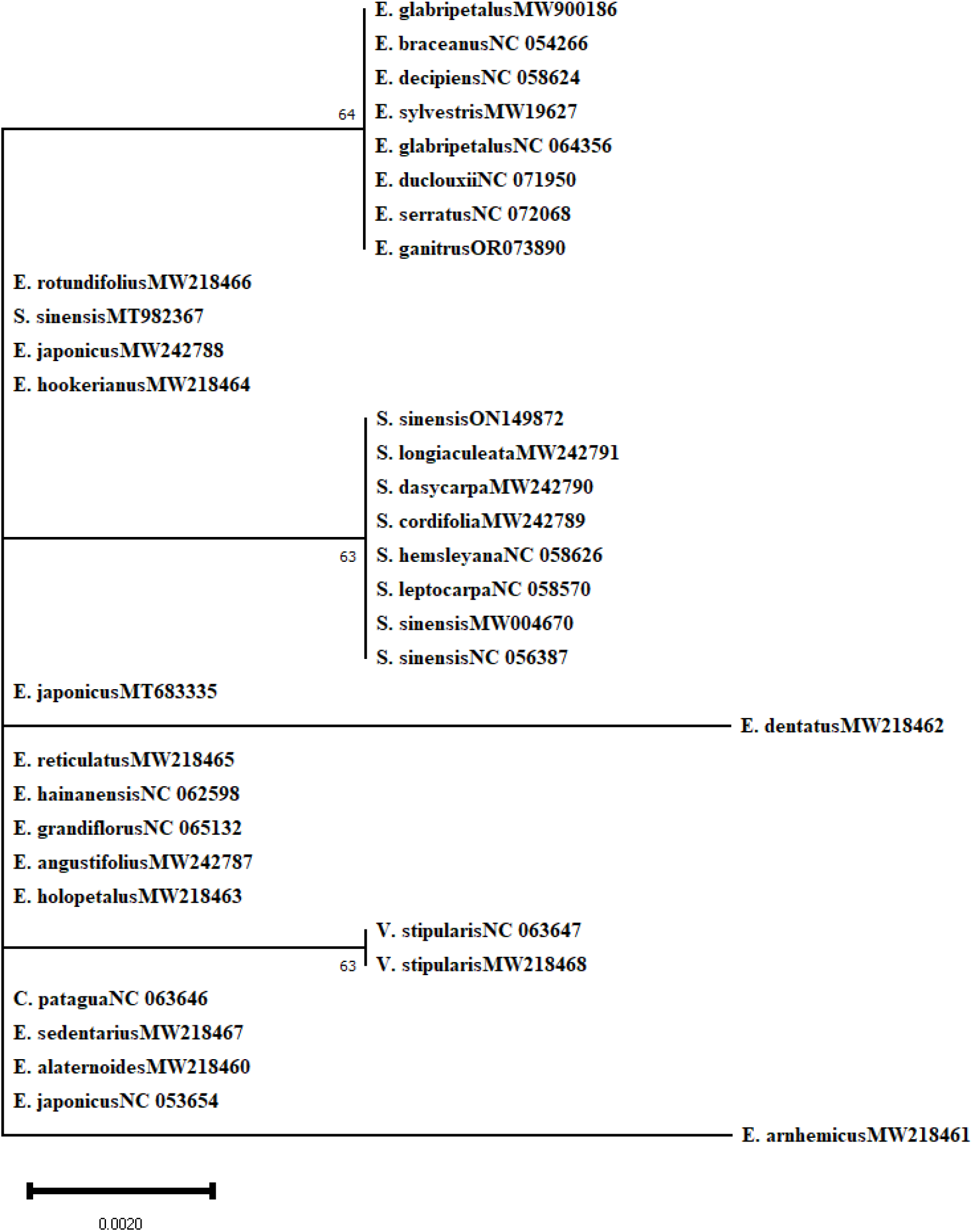
Neighbor-joining tree of Elaeocarpaceae based on *accD* sequences using Tamura 3-parameter method. This analysis involved 34 nucleotide sequences and 254 sites. The Number on the branches represent more than or equal to 50 percent support after the 10,000 bootstrap replications test.

The concatenated sequences were assembled, and phylogenetic relationships based on the Neighbor-Joining (NJ) method were established using *ndhJ*+*matK*, *rpoB*+*matK*, and *accD*+*matK* sequences for twenty-seven plants of Elaeocarpaceae family (Fig. 8-10). According to the phylogenetic tree generated using two-locus barcodes, all 19 *Elaeocarpus* and five *Sloanea* species (except *S. sinensis,* MT982367) were clustered with strong bootstrap support. The phylogenetic analysis showed that single (*matK*, *rpoB ndhJ*, *accD*) and the combined (*ndhJ*+*matK*, *rpoB*+*matK*, and *accD*+*matK)* chloroplast sequences exhibit better identification ability at the genus level, indicating their applicability in the identification of Elaeocarpaceae plants. However, the Neighbor-Joining tree based on *accD* loci was poorly resolved, and some taxa of the genus *Ealeocarpus* were poorly separated.

**Fig. 8.**
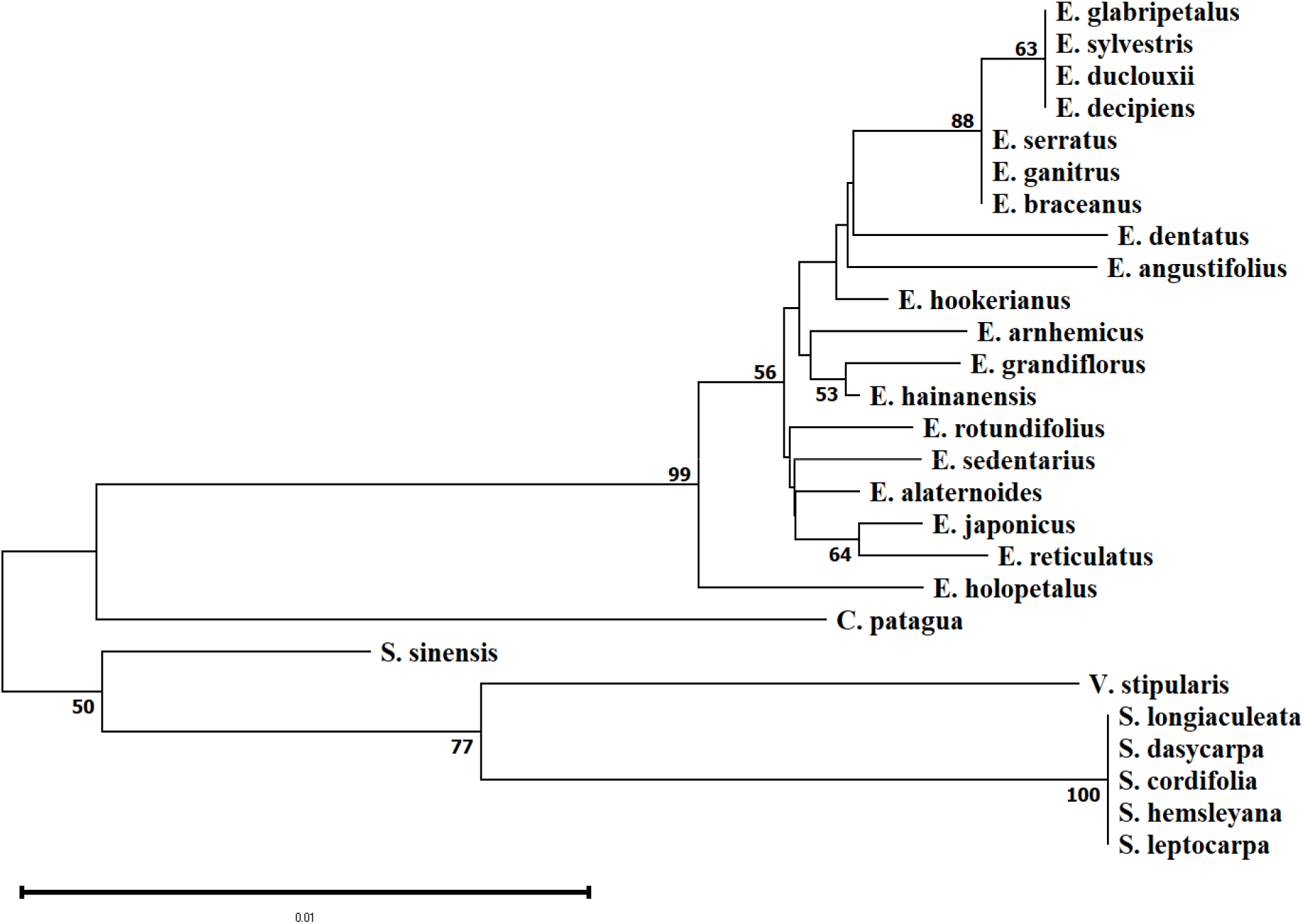
Neighbor-joining tree of Elaeocarpaceae based on *rpoB*+*matK* sequences using Tamura 3-parameter method. This analysis involved 27 nucleotide sequences and 937 sites. The Number on the branches represent more than or equal to 50 percent support after the 10,000 bootstrap replications test.

**Fig. 9.**
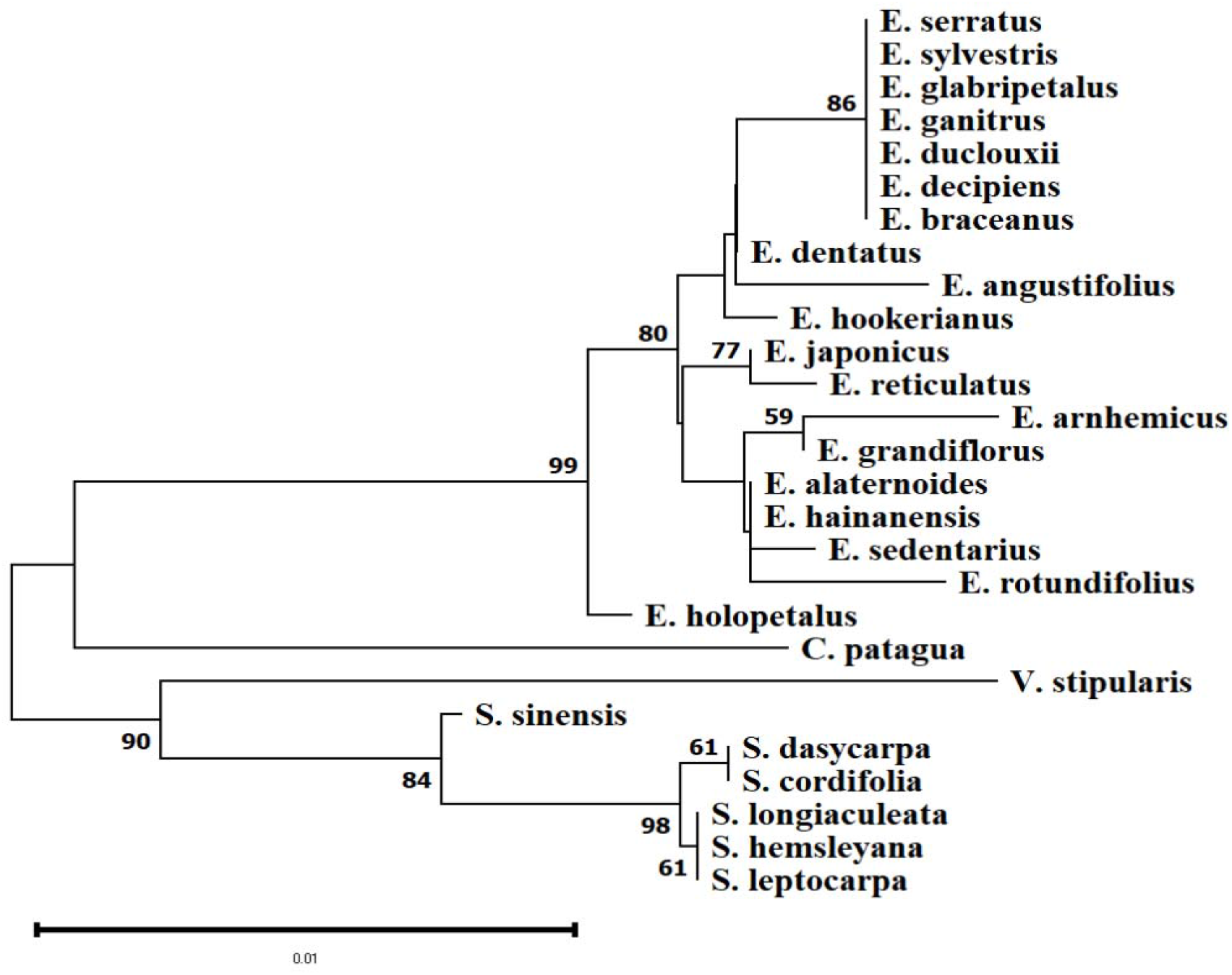
Neighbor-joining tree of Elaeocarpaceae based on *ndhJ*+*matK* sequences using Tamura 3-parameter method. This analysis involved 27 nucleotide sequences and 872 sites. The Number on the branches represent more than or equal to 50 percent support after the 10,000 bootstrap replications test.

**Fig. 10.**
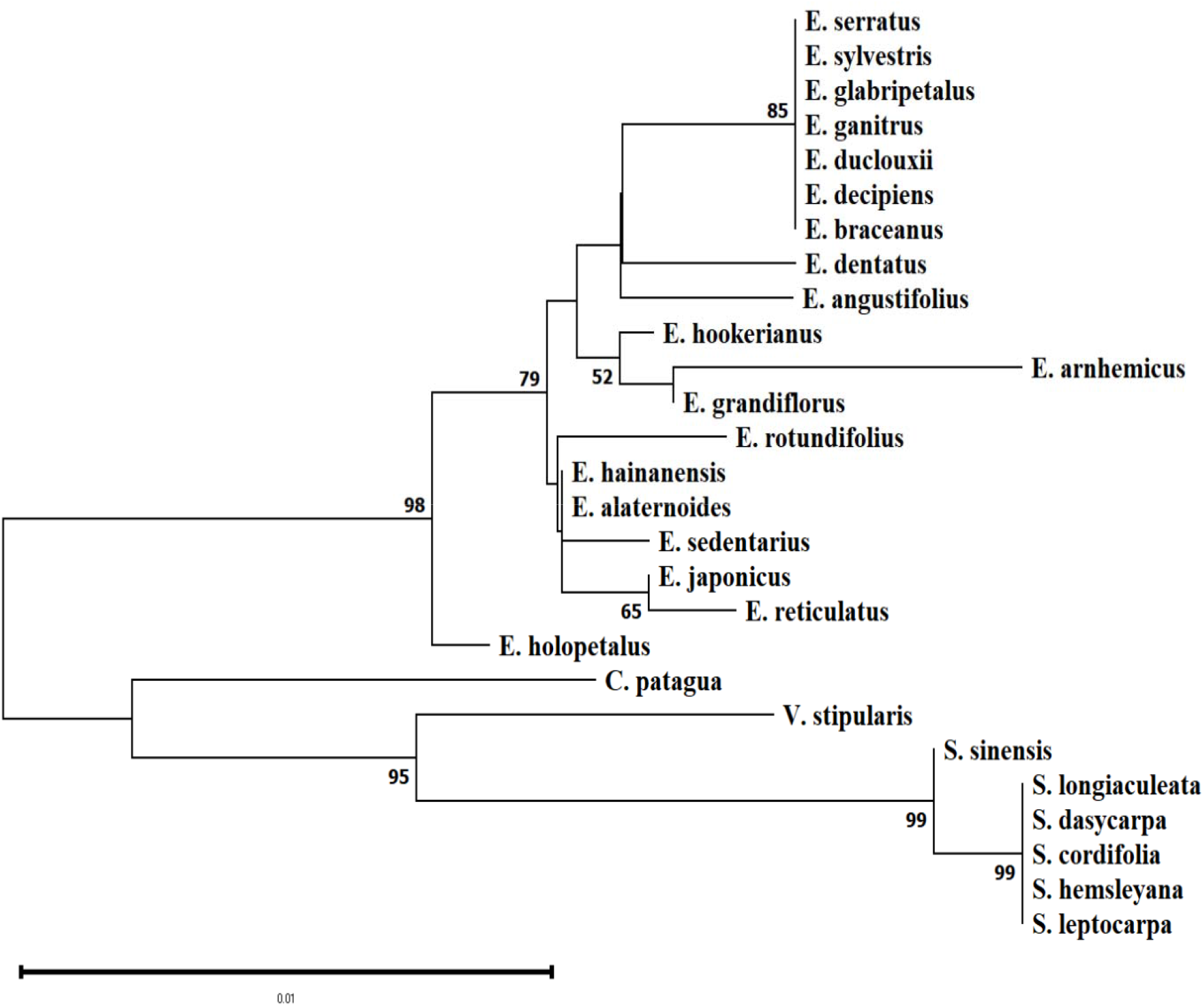
Neighbor-joining tree of Elaeocarpaceae based on *accD*+*matK* sequences using Tamura 3-parameter method. This analysis involved 27 nucleotide sequences and 649 sites. The Number on the branches represent more than or equal to 50 percent support after the 10,000 bootstrap replications test.

### DNA Barcoding gap analysis and Species delimitation

An ideal DNA barcode should exhibit greater significant inter-specific than intra-specific genetic variance. In order to evaluate the potential of single (*matK*, *rpoB ndhJ*, *accD*) and combined barcode sequences (*ndhJ*+*matK*, *rpoB*+*matK*, *accD*+*matK*) in the development of species-specific DNA barcodes, the barcoding gap analysis was carried out based on frequency distribution histogram and pairwise distance divergence using Automatic Barcode Gap Discovery server. In the *matK* and *ndhJ*+*matK* sequences, the maximum intra-specific genetic distances were significantly smaller than the minimum inter-specific genetic distances, thus forming a prominent barcoding gap. The barcoding gap analysis of other barcode sequences (*rpoB ndhJ*, *accD*, *rpoB*+*matK*, *accD*+*matK*) sequences produced no distinct barcoding gap, which might be due to overlapping at intra- and inter-specific distances or less number of taxon (Fig. 11).

**Fig. 11.**
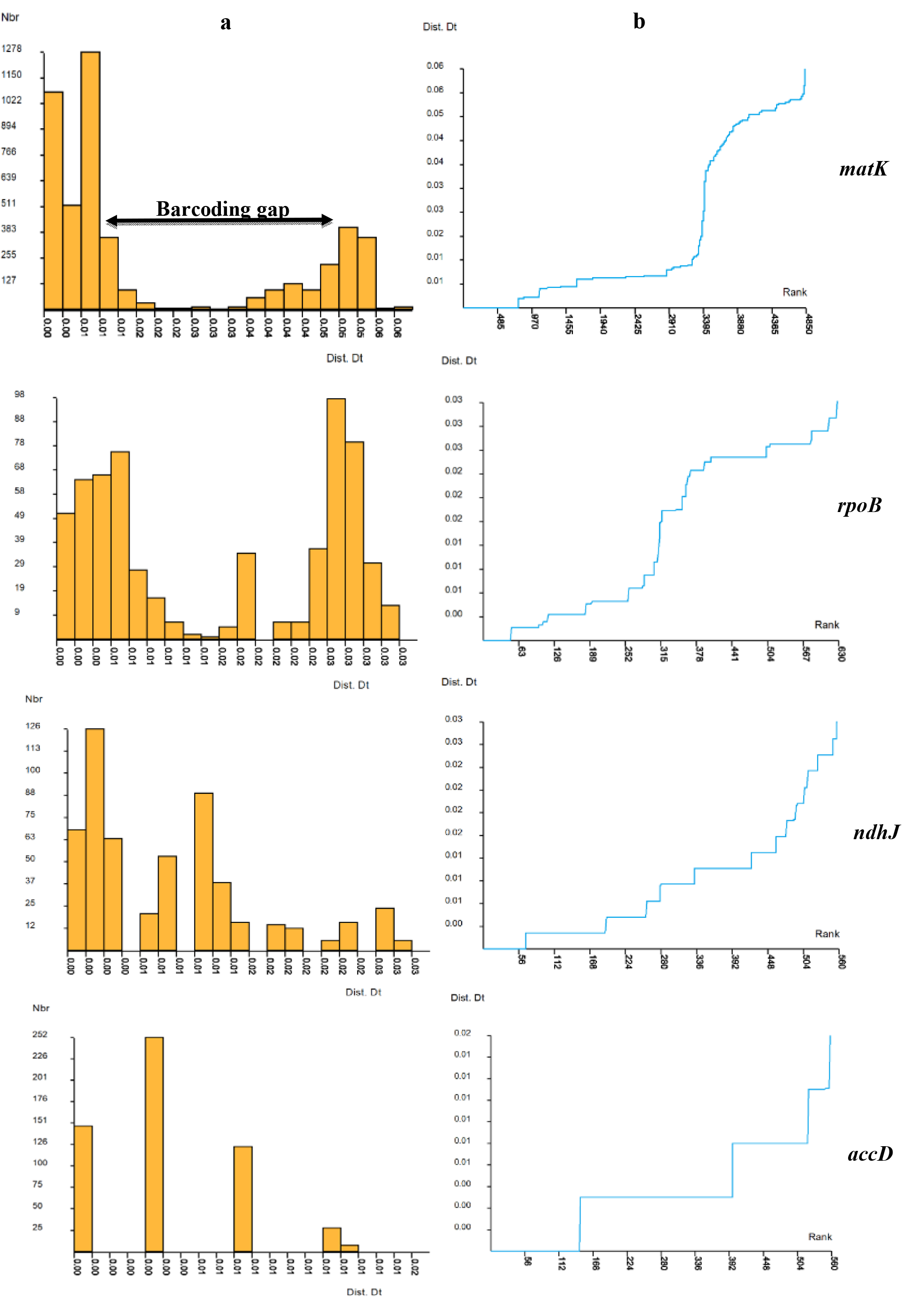
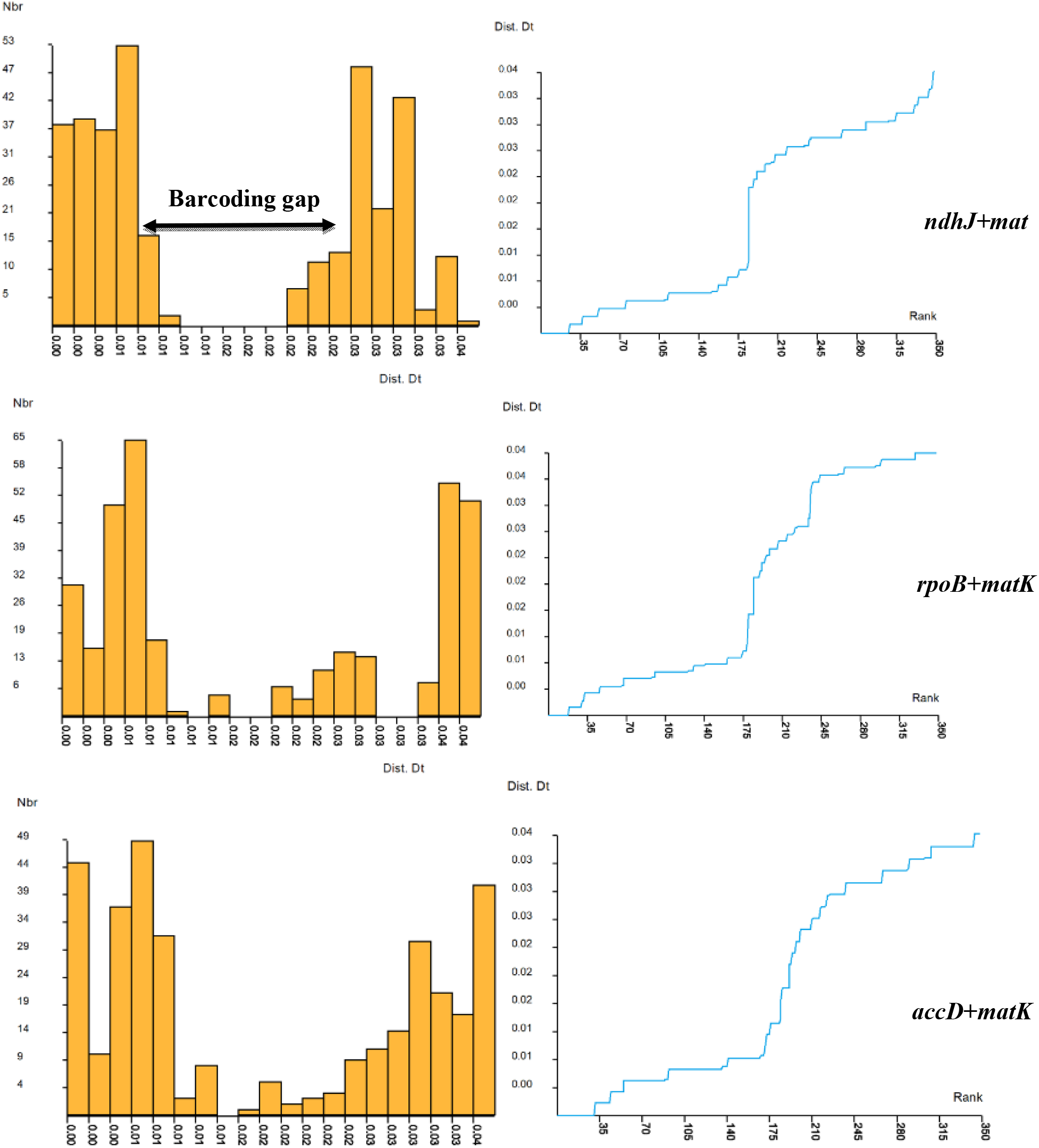
Results of Assemble Species by Automatic Partitioning (ASAP) analysis for candidate barcode sequences. (a) Distribution of pairwise differences, (b) Ranked pairwise differences.

Each sequence set (*matK*, *rpoB*, *ndhJ*, *accD*, *rpoB*+*matK*, *ndhJ+matK*, *accD*+*matK*) was partitioned into subsets independently using the ASAP (Assemble Species by Automatic Partitioning) algorithm based on pairwise genetic distances. The number of Operational Taxonomic Units (OTUs), W-values (Ranked distance), and ASAP score were measured for each partition. Partitions with the lowest ASAP score are considered the best [27] (Table 6). The sequence partition with the lowest ASAP score (3.50) distributed the sequences into 4 OTUs and the threshold dist. value 7.66e-05 based on the *ndhJ* gene sequence. Similarly, the sequence partition with the lowest ASAP score (1.00) distributed the sequences into 4 OTUs and the threshold dist. value 1.38e-03 based on the *rpoB* gene sequence. Based on the *accD* gene, the sequence partition with the lowest ASAP score (2.00) distributed the sequences into 3 OTUs and the threshold dist. value 0.005931. The sequence partition with the lowest ASAP score (1.5) distributed the sequences into 3 OTUs and the threshold dist. value 0.025844 based on *matK* gene sequence. The sequence partition with the two-locus barcode (*ndhJ+matK*) generated the lowest ASAP score (1.00) and distributed the sequences into 3 OTUs and threshold dist. value 0.020992.

The sequence partition with the lowest ASAP score (1.00) distributed the sequences into 5 OTUs and the threshold dist. value 0.010817 based on the *rpoB+matK* gene sequence. Based on *accD+matK*, the sequence partition with the lowest ASAP score (1.50) distributed the sequences into 4 OTUs and the threshold dist. value 0.010887. The partition analysis based on *ndhJ*, *rpoB*, and *accD+matK* gene sequences demonstrated the clustering pattern of species at the genus level.

### Species-specific barcodes based on Single Nucleotide Polymorphisms (SNPs)

Species-specific barcodes were developed by conducting a Single-nucleotide polymorphism (SNP) analysis of each candidate barcode sequence alignment (*matK*, *rpoB, ndhJ,* and *accD*), and the identified sequence was blasted against the NCBI nucleotide database for further authentication. Specific barcodes were generated for *Elaeocarpus arnhemicus, Elaeocarpus japonicus, Elaeocarpus sericopetalus, Elaeocarpus rotundifolius,* and *Elaeocarpus reticulatus* based on *matK* gene. Information about specific barcodes of *Elaeocarpus reticulatus, Elaeocarpus holopetalus, Elaeocarpus alaternoides, Elaeocarpus grandiflorus, Elaeocarpus dentatus, Elaeocarpus alternifolius,* and *Elaeocarpus angustifolius* was obtained based on the *rpoB* gene. Based on the *ndhJ* gene, the specific barcode of the species *Elaeocarpus rotundifolius, Elaeocarpus arnhemicus, Elaeocarpus angustifolius*, and *Elaeocarpus glabripetalus* were obtained. Similarly, specific barcodes for *Elaeocarpus arnhemicus* and *Elaeocarpus dentatus* were generated based on the *accD* gene. A combinational barcode analysis produced more barcodes than a single locus due to more mutational and single nucleotide polymorphism sites. However, no specific barcodes for genera *Crinodendron, Sloanea,* and *Vallea* were observed, possibly due to insufficient species within the genus. The two-dimensional code of each specific barcode was created, which can be used for species identification and easily scanned by electronic equipment. The specific barcodes provide a foundation for future DNA barcoding and germplasm conservation for the *Elaeocarpus* gen*us* (Fig. 12; Table 7).

**Fig. 12.**
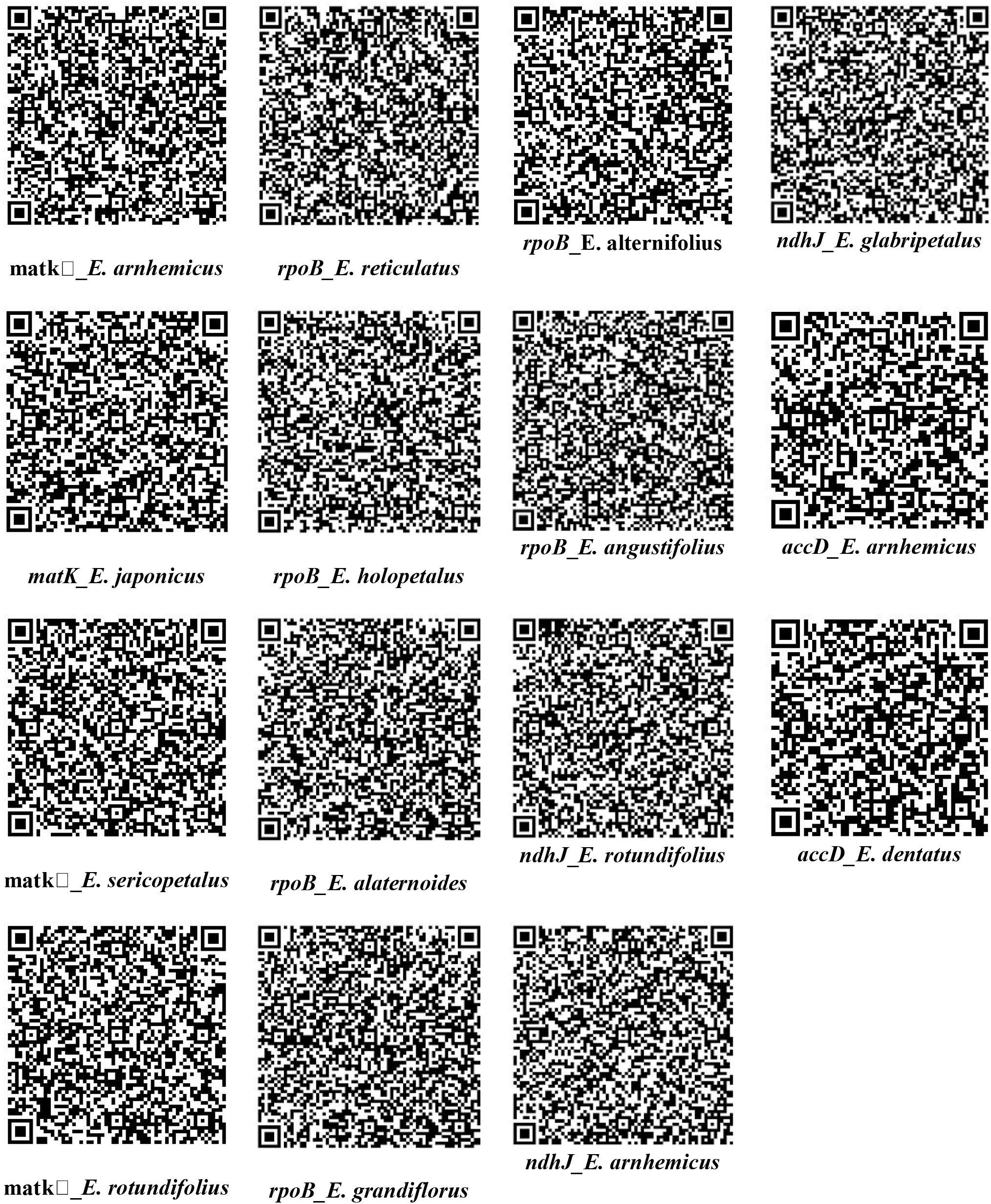
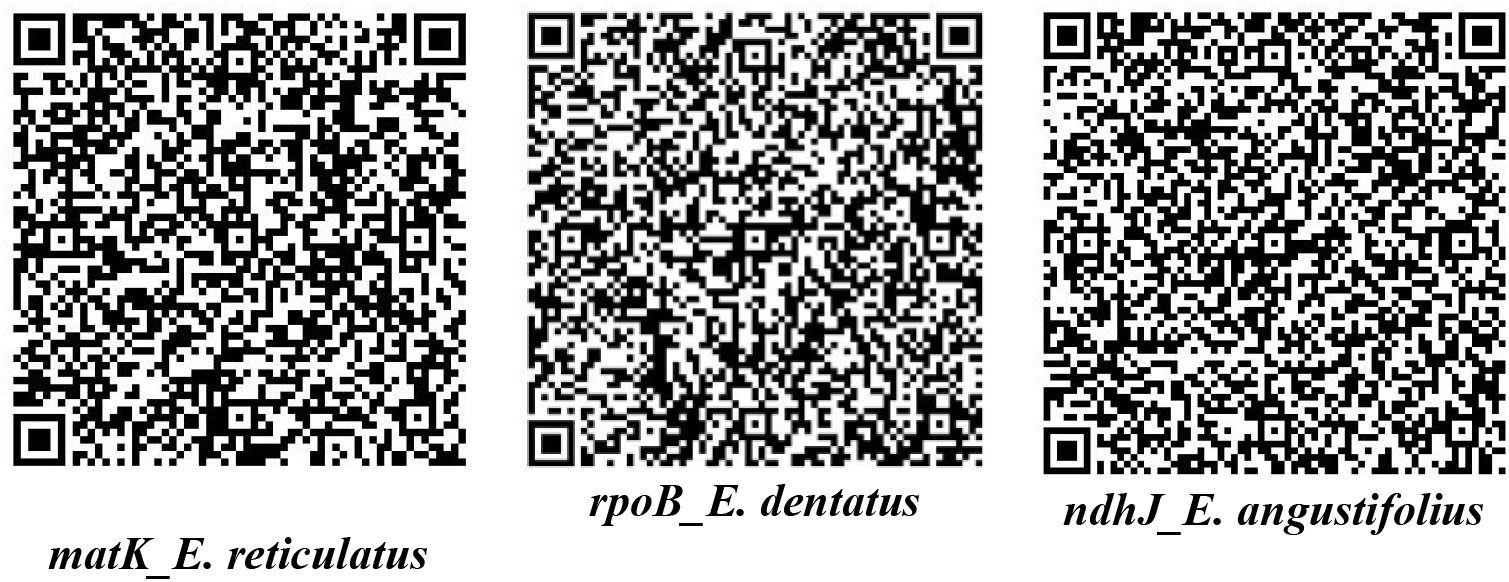
Two-dimensional DNA barcodes developed for Elaeocarpaceae plants based on single nucleotide polymorphisms analysis of each candidate barcode sequence alignment.

## Conclusion

The application of chloroplast-based DNA barcodes (*matK*, *rpoB, ndhJ, accD, ndhJ+matK, rpoB+matK,* and *accD+matK*) in identifying Elaeocarpaceae plants was confirmed by *In silico* analysis. Polymorphism site analysis indicated that *matK* and *rpoB+matK* sequences had the highest proportion of variable and parsimony informative sites. Phylogenetic analysis using the single (*matK*, *rpoB, ndhJ,* and *accD)* and combined (*ndhJ*+*matK*, *rpoB*+*matK*, and *accD*+*matK*) sequences is nearly consistent. Neighbor-joining analysis indicated that the discrimination capacity of *matK*, *rpoB, ndhJ, accD*, *ndhJ*+*matK*, *rpoB*+*matK*, and *accD*+*matK* loci are variable among Elaeocarpaceae plants, in which *matK, rpoB,* and ndhJ loci reveal as a more potential locus for DNA barcoding. The barcoding gap analysis of barcode sequences resulted in a prominent barcoding gap produced by *matK* and *ndhJ*+*matK* loci. However, *rpoB, ndhJ*, *accD*, *rpoB+matK*, and *accD+matK* sequences produced no distinct barcoding gap, possibly due to overlapping at intra- and inter-specific distances or less taxon numbers. The partition analysis based on *ndhJ*, *rpoB*, and *accD+matK* gene sequences demonstrated the exact clustering pattern of species within the genus. Single Nucleotide Polymorphisms (SNP) are one of the best measures of phylogenetic diversity because they can distinguish closely related species. The species-specific DNA barcodes for the genus *Elaeocarpus* were generated using the SNP sites. The study also expands the application of plastid-based barcodes in DNA barcoding and offers an innovative approach for plant identification at the species level. The present study provides insight into understanding the phylogenetic relationship of Elaeocarpaceae plants and could be helpful in the germplasm conservation and utilization programs of the genus *Elaeocarpus*.

